# Integrative Computational Framework for Understanding Metabolic Modulation in Leishmania

**DOI:** 10.1101/512277

**Authors:** Nutan Chauhan, Shailza Singh

## Abstract

The integration of computational and mathematical approaches is used to provide a key insight into the biological systems. Here, we seek to find detailed and more robust information on *Leishmanial* metabolic network by performing mathematical characterization in terms of Forman/Forman-Ricci curvature measures combined with flux balance analysis (FBA). The model prototype developed largely depends on its structure and topological components. The correlation of curvature measures with various network statistical properties revealed the structural-functional framework. The analyses helped us to identify the importance of several nodes and detect sub-networks. Our results revealed several key high curvature nodes (metabolites) belonging to common yet crucial metabolic, thus, maintaining the integrity of the network which signifies its robustness. Further analysis revealed the presence of some of these metabolites in redox metabolism of the parasite. MGO, an important node, has highly cytotoxic and mutagenic nature that can irreversibly modify DNA, proteins and enzymes, making them nonfunctional, leading to the formation of AGEs and MGO^●-^. Being a component in the glyoxalase pathway, we further attempted to study the outcome of the deletion of the key enzyme (GLOI) mainly involved in the neutralization of MGO by utilizing FBA. The model and the objective function both kept as simple as possible, demonstrated an interesting emergent behavior. The nonfunctional GLOI in the model contributed to ‘zero’ flux which signifies the key role of GLOI as a rate limiting enzyme. This has led to several fold increase production of MGO, thereby, causing an increased level of MGO^●-^ generation. Hence, the integrated computational approaches has deciphered GLOI as a potential target both from curvature measures as well as FBA which could further be explored for kinetic modeling by implying various redox-dependent constraints on the model. Designing various *in vitro* experimental perspectives could churn the therapeutic importance of GLOI.

**Author Summary:** Leishmaniasis, one of the most neglected tropical diseases in the world, is of primary concern due to the increased risk of emerging drug resistance. To design novel drugs and search effective molecular drug targets with therapeutic importance, it is important to decipher the relation among the components responsible for leishmanial parasite survival inside the host cell at the metabolic level. Here, we have attempted to get an insight in the leishmanial metabolic network and predict the importance of key metabolites by applying mathematical characterization in terms of curvature measures and flux balance analysis (FBA). Our results identified several metabolites playing significant role in parasite’s redox homeostasis. Among these MGO (methylglyoxal) caught our interest due to its highly toxic and reactive nature of irreversibly modifying DNA and proteins. FBA results helped us to look into the important role of GLOI (Glyoxalase I), the enzyme that catalyses the detoxification of MGO, in the pathway that, when non-functional, has resulted into increased level production of free radicals and AGEs (advanced glycation end products). Thus, our study has deciphered GLOI as a potential target which could further be explored for future *in vitro* experiments to design potential GLOI inhibitors.

## 1. Introduction

Networks (graphs) are an important part of our everyday life. Network science deals with different complex networks, *viz*., social networks, telecommunication networks, biological networks; according to the types of nodes (vertices) communicating with each other with the help of edges (links). In biology, networks have always been present in many forms, such as, brain network, immune network, genetic linkage maps, signaling network, protein-protein interaction network, reaction network, metabolic networks, etc. Among them, metabolic networks are becoming of much interest amongst the researchers due to their unique feature of connecting every conserved node (the metabolites) with the links (the reactions) that is catalyzed by specific gene products responsible for growth and maintenance of a cell [1].

Moreover, metabolic networks can be used to shed light on finding various disease mechanisms via the identification of essential genes by implying perturbations on networks, thereby, characterizing the structure-function relationships [2-8]. Structure-function relationship of a network in terms of network *robustness* and reliability is strongly linked to its geometric and topological properties. In biology, the network robustness is defined as the ability of network to withstand induced perturbances that helps to understand the nature of diseases and to recover from it. To study the structure-function relationships and geometrical characterization of these models, there are several ways for quantitatively analyzing complex networks using tools derived from mathematical aspects of graph theory [8]. To unravel the meaningful information of any complex network, network analysis helps in reducing the system to create a basic structural framework as a means for understanding the relation between the elements and their interactions. Recent technological advances in graph theory have allowed us to analyze and describe highly complex systems at smaller and more detailed scales. Application of graph theory have been utilized to study global impact of long term sickle cell disease on brain [9], to diagnose pre-symptomatic Alzheimer’s disease [10-12], to understand the human brain network [13,14]. Recently, the development of geometry based measures is significantly attracting the focus of researchers to characterize the structural aspect of complex networks [15-26].

### 1.1. Curvature-based methods for complex networks

In geometry based measures of complex networks *curvature* is one of the notions that are being explored to understand the complexity of graphs. Curvature plays central role in Riemannian geometry since it represents a measure to quantify the deviation of a geometrical object from being flat [27]. Curvature measures can be used to quantify the robustness and thereby the functionality of networks.

There are several types of curvature notions, among them *Ricci curvature* is known to be the most useful for analyzing the complex networks [16-23, 28-33]. Ricci curvature measures deviance of geodesics (shortest path) relative to Euclidean shortest-paths and is related to mass transport or entropy (Wasserstein metrics) [34-36]. Basically, Ricci-curvature measures the related mass transport and entropy, hence, a higher value of Ricci curvature is directly related to the higher mass/entropy. High Ricci curvature is typically found near network hubs, where an anchor node (high curvature node) is connected to many existing nodes in close proximity. Principally, curvature is intimately connected to network entropy, in turn, is very close to robustness. Therefore, higher curvature on a node/edge suggests that removal of high curvature node/edge from the network will cause loss of a lot of mass, consequently the network will collapse.

There are two different discretization of Ricci curvature *Ollivier-Ricci curvature* [16,33] and *Forman-Ricci curvature* [37]. These two curvature notions are seen to be particularly appealing and give efficient solutions for network geometrizations. In case of undirected networks, Ollivier-Ricci curvature has proved its applicability in various network analyses [17-20,22,23,38,39]. On the other hand, Forman-Ricci curvature has been introduced as a tool for undirected [29,40] as well as directed [41] network analyses. Irrespective of their applicability and handling the large networks, the two curvature notions demonstrated high correlation [42]. However, researchers suggested that Forman-Ricci curvature is a faster computation method and can be utilized in larger real-networks to gain preliminary insight into Ollivier-Ricci curvature that is much more computationally extensive. Additionally, it was shown that Forman-Ricci curvature can handle weights of edges as well as of vertices in contrast to Ollivier-Ricci curvature, where only weighted edges are analyzed with an additional step of normalization of neighboring edge weights. This property of Forman curvature makes the measure suitable for the analysis of both unweighted and weighted networks [29]. Current study exploits the application of Forman-Ricci curvature and systems biology to capture the behavior of Leishmanial metabolic network and describe the interplay between small and large scale networks.

### 1.2. Leishmaniasis and systems-biology

Neglected infectious diseases have affected at least a billion of human populations worldwide [43, 44] and are of primary concern due to the lack of effective and affordable drug regimens. Among them, cutaneous leishmaniasis (CL), a very common clinical form of leishmaniases, has always been overlooked as a major public health problem due to its non-fatality. The severity of the disease ranges from disfigurement and residual scars. The causative agent of CL, a protozoan parasite, *Leishmania major* has a digenetic lifecycle and lives in two hosts, sandfly, *Phlebotomus argentipes*, and human, in the form of flagellated promastigotes (procyclic phase) and non-flagellated amastigotes (metacyclic phase), respectively.

When inside the human host, the parasite has to undergo the oxidative stress generated by host macrophages (Figure 1). This stress is also attained from free radical generation within the parasite in diverse ways. This mainly includes electron transport chain where superoxide free radicals are consistently generated. Moreover, there are several highly reactive metabolites such as glyoxals that contribute to the generation of free radicals. These sorts of metabolites can react with other metabolites and enzymes. This not only causes inactivation of enzyme activity but also contribute in the formation of free radicals like super oxide anion and peroxynitrite, etc. One such highly reactive metabolite, methylglyoxal (MGO), is the byproduct of glycolysis cycle, where the triose phosphate intermediates dihydroxyacetone phosphate (DHAP) and glyceraldehyde 3-phosphate (GA3P) eliminate phosphate via an enediolate intermediate [45]. While inside the macrophage, leishmania demands high energy because of the rapid proliferation rate that results in high rates of glycolysis, thereby, increased production of MGO. Also, there are few minor sources of methylglyoxal production which includes aminoacetone and hydroxyacetone, intermediates generated during catabolism of threonine and acetone [45].

**Figure 1.**
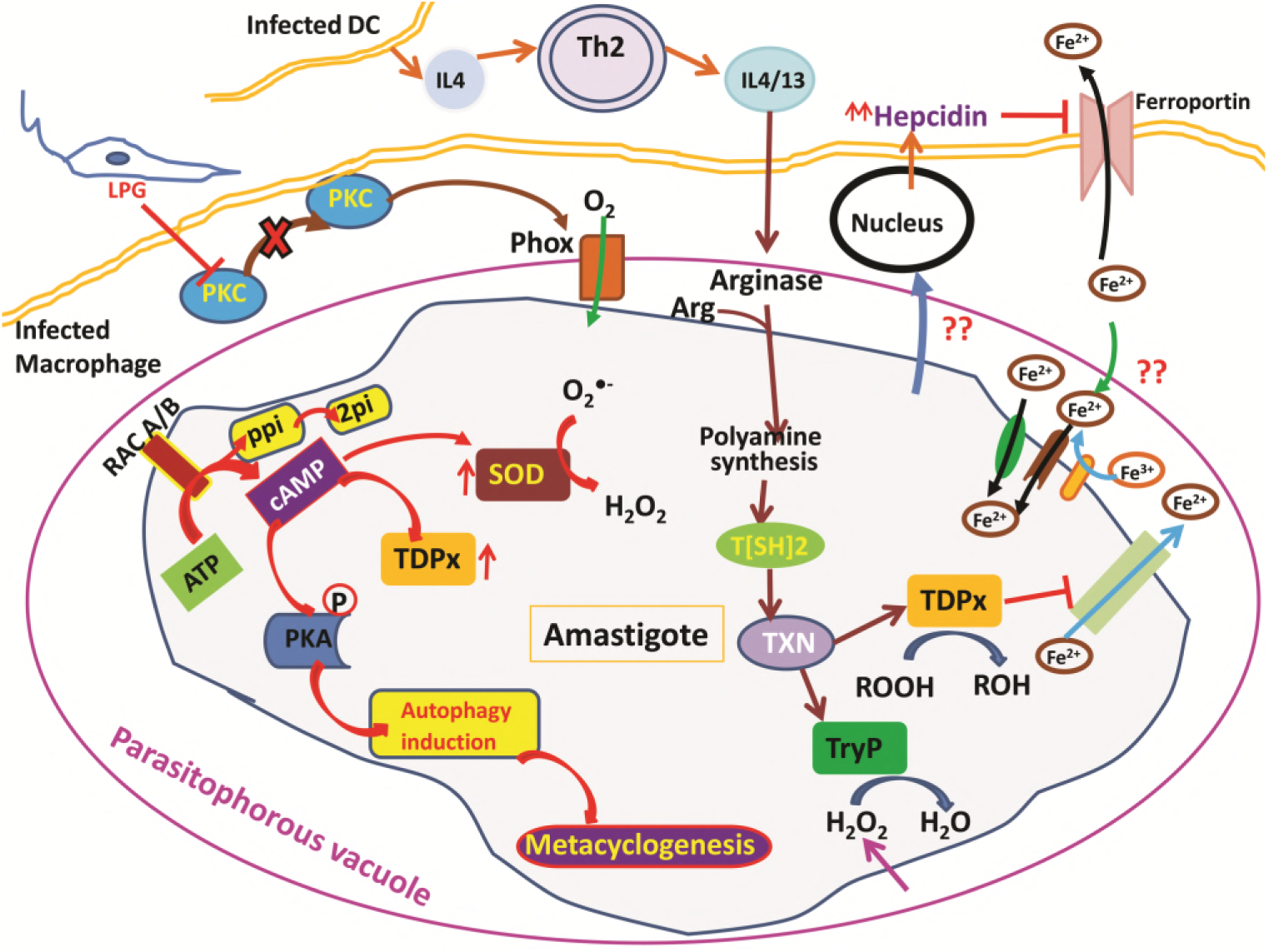
Illustration of *Leishmania* parasite survival strategies from oxidative stress

To deal with these types of internal and external trauma and to survive inside the lethal lysosomal environment of macrophage by escaping from reactive nitrogen and oxygen species, the parasite has evolved with its own unique redox machinery that helps in neutralization of free radicals. Leishmania parasite possess an exceptional oxidant and chemical defense mechanism, involving a very unique small molecular weight dithiol, trypanothione (T[SH]_2_), that helps the parasite to manage its survival inside the host macrophage. The main component of this antioxidant defense mechanism on which the parasite relies on are trypanothione synthase (TryS), trypanothione reductase (TryR), tryparedoxn peroxidase (TryP), tryparedoxine-dependent peroxidase (TDPx), and tryparedoxin (TXN). The center of these components is a central reductant, T[SH]_2_ that is synthesized by TryS enzyme. The reduced state of T[SH]_2_ is maintained by TryR enzyme that is an functional analogue of human glutathione reductase (GR) [46, 47]. T[SH]_2_ helps in reduction of TXN that further reduces TryP and TDPx for detoxification of reactive species generated during synthesis of hydroperoxide, deoxyribonucleotide, and reactive nature of methylglyoxal. Capability of T[SH]_2_ of reducing free reactive species is 600 times faster rate than GR mechanism of human host which makes the parasite redox metabolism unique.

Due to the incapability and increasing drug resistance of the current therapy regime techniques (chemotherapy) against leishmaniasis, the necessity to develop novel, more efficient and affordable anti-leishmanial drugs is desirable. Moreover, more effective molecular drug targets with therapeutic value are also needed. The difference between *Leishmania* and human host redox metabolism attracts researcher’s interest to explore this mechanism for discovering novel drug targets and design potential inhibitors against the redox enzymes without crossing the host machinery [48].

Redox metabolism consists of main redox components as well as other connecting molecules like enzymes, signaling molecules, etc. To study the role and effect of other intermediates in the redox network, systems biology approaches including computational and quantitative modeling methods are nowadays being used [49]. Systems biology approaches have already been utilized to study systems dynamics of complex redox metabolic pathways in many organisms [50], such as, *Escherichia coli*, purple non-sulfur bacteria [51], *Saccharomyces cerevisiae* [52-54]. Redox systems have been characterized from systems biology perspective using top-down and bottom-up approaches. Moreover, omics data is also being integrated to construct metabolic kinetic models to explore the complex mechanisms [52, 55]. A wide spectrum of techniques applied for analysis of biochemical networks include genome scale metabolic modeling (GEMs) [50, 52-54], constraint based methods, kinetic pathway modeling [56], interaction-based analyses, Petri nets, etc [57-59]. Before implementing any of these techniques, the knowledge and availability of all possible components and their inter-connections is a pre-requisite. Recently, kinetic modeling and constraint-based metabolic modeling have gained researcher’s focus. In case of kinetic modeling, the availability of kinetic data and rate laws is required for simulating the network by using first order *ordinary differential equations* (ODE). Kinetics of several bacteria to study the antibiotic resistance and gene transfer [60,61] has been studied using kinetic modeling. However, due to the lack of sufficient kinetic data the use of metabolic kinetic modeling on larger scale networks is limited. On the other hand, constraint-based analysis such as *flux-based analysis (FBA)* requires only the knowledge of stoichiometry of the metabolites and can easily be applied to even larger scale metabolic networks.

### 1.3. Flux balance analysis(FBA)

Metabolic modeling using FBA has proved to be effective in numerous ways such as in identifying metabolic gaps and blocked reactions to get an insight in finding newer drug therapy. FBA calculations rely on the assumption of steady-state growth and mass balance and can be used to predict the flux of a metabolite flowing through a metabolic network. This property of FBA makes it possible to predict the growth rate of an organism under several applied constraints. To predict the growth rate or to find a specific cellular function, FBA tries to find possible solutions to optimize the stated objective function(s) and gives quantitative insights into the genotype-phenotype relationship of the model. Although, FBA is incapable of predicting the concentrations of metabolites and does not account for any regulatory effects like activation/inactivation of enzymes or gene-expression regulations, there is wide range of literature showing its applications in many fields [62]. FBA has been explored in many organisms such as drug-sensitive and resistant *Plasmodium falciparum* [63] was studied to predict chloroquine mediated dose dependent DNA inhibition, *Leishmania infantum* and *Leishmania donovani* [64,65.] to identify important anti-leishmanial drug targets. In *P. falciparum* metabolic network gene-knockout experiment helped to spot the importance of thioredoxin reductase through FBA [66]. Therapeutic importance of promising drug targets in *L. major* [67], *P. falciparum* [68, 69], *T. cruzi* [70] and in many more metabolic networks through FBA mediated metabolic analysis [71-75].

In current work, we attempted to explore the application of *Forman* and *Forman-Ricci curvature* on metabolic network of *Leishmania*. Using curvature measures, we tried to identify important metabolic pathways contributing to the overall network robustness by recognizing “important” metabolic hubs in the network. Through this, we seek to churn metabolites and enzymes having essential role in the survival of the parasite. Further, to predict the effect of the absence of these “important” metabolites, constraint-based FBA is used to uncover the importance of enzymes and their probable role as drug target of therapeutic significance (Figure 2).

**Figure 2.**
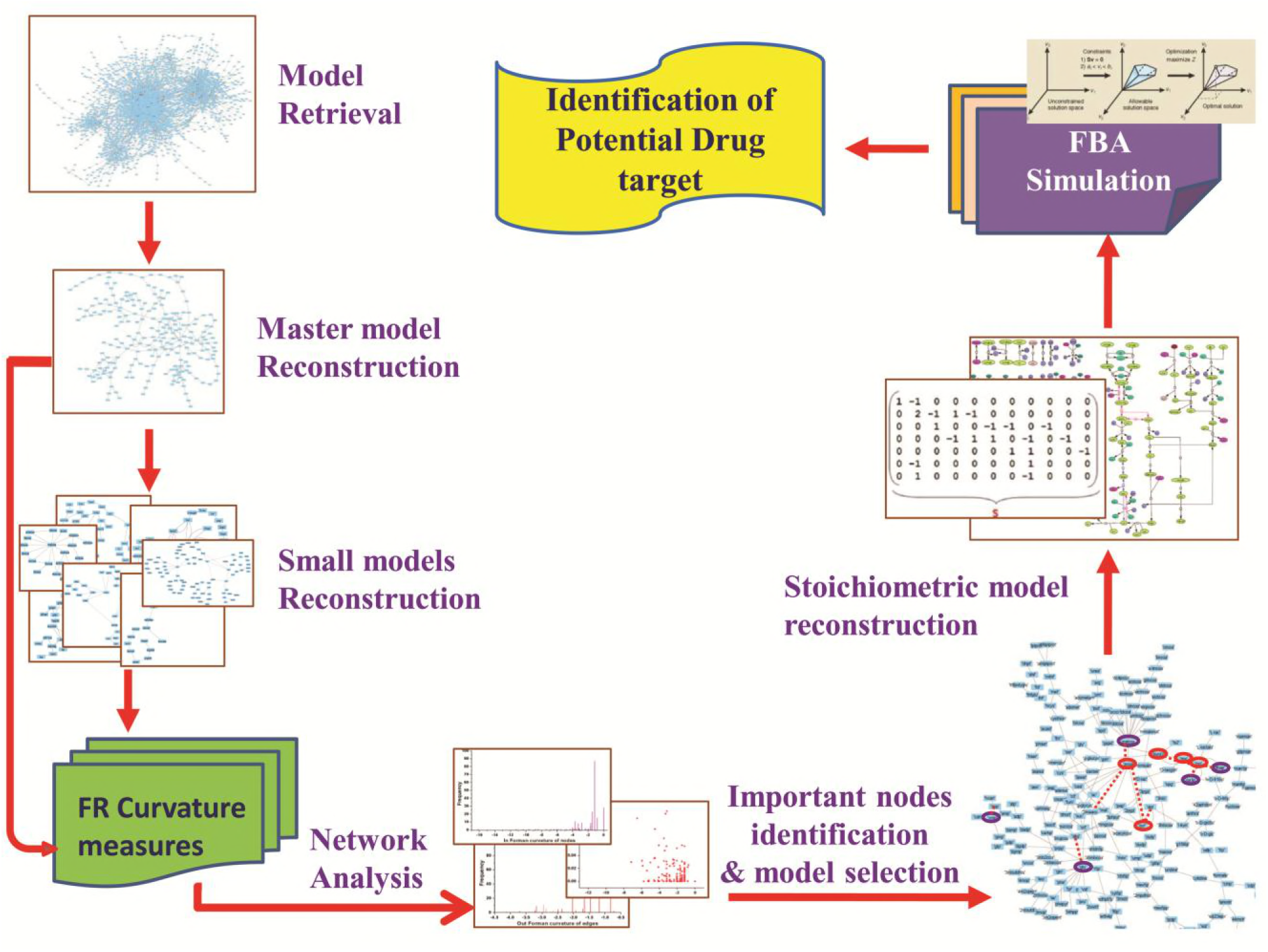
Overview of the workflow strategy adopted

## 2. Material and Methodology

### 2.1 Data sets and model reconstruction

Initially, *Leishmania major* metabolic network **iAC560** [76] was retrieved from BioModels Database and imported to Cell Designer (4.4) [77]. The model consisted of 1112 reactions and 1147 metabolites. This model is used to construct a bipartite directed graph and considered as master model. The master model is further used for constructing small scale models (Figure 2).

### 2.2 Generation of connectivity matrix

Connectivity matrices for master and other models were generated by considering the connections between metabolites (substrate to product). A value of ‘1’ is assigned for connection, if a metabolite (substrate) is connected (producing or converting into) to other metabolite (product), and ‘0’ for no connection.

### 2.3 Network analysis

#### 2.3.1 Network Topology

The reconstructed bipartite directed graphs were visualized in Cytoscape (3.6.0) [78] and their topological information (network properties) of nodes and edges were obtained using graph theory. Various properties including degree of nodes, betweenness centrality (the number of shortest paths that passes through a node), closeness centrality (the measure of mean distance from a node to other nodes), clustering coefficient (a measure of the degree to which nodes in a graph tend to cluster together), etc. were calculated in NetworkAnalyzer module.

In geometry based measures of complex networks curvature is used to quantify the deviation of an object from being a flat plane. Among the several curvature measures, we used Forman curvature and Forman-Ricci curvature for the analysis of our bipartite directed graphs. For any given directed bipartite graph we calculated [40]:

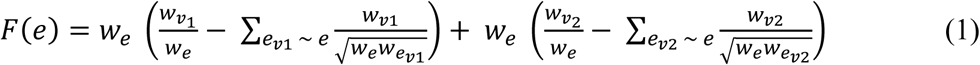

Where, *F* is the Forman curvature of the directed edge 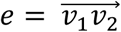 that originates from node *v_1_* and terminates at node *v_2_*. Only those directed edges were considered form calculation that either terminate at node *v_1_* or originate at node *v_2_*. Furthermore, self-loops or self-edges on nodes *v_1_* and *v_2_*, edges facing opposite direction, were completely ignored.

##### 2.3.2.1 Forman curvature on Networks

To distinguish between incoming, *E_I,v_*, and outgoing, *E_O,v_*, edges for a given node *v*, unnormalized Forman curvature [79] (the sum of the curvature of all edges incident or outgoing on that node), *In Forman curvature **F_I(v)_*** and *Out Forman curvature **F_O(v)_*** were computed as follows:

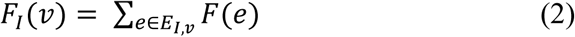

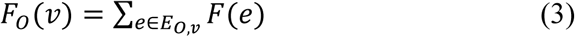

And, the total flow on a given node was obtained as:

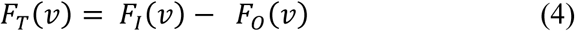

##### 2.3.2.2 Forman-Ricci curvature on Networks

Moreover, for determining Forman-Ricci curvature, *FR(v)* (normalized Forman curvature) [79] of a node *v*, the sum of the curvature of all edges incident or outgoing on that node was divided by the degree of that node:

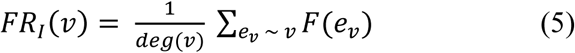

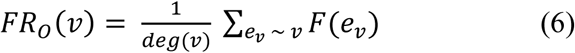

Where, *FR_I(v)_* and *FR_O(v)_* are *In* and *out Forman-Ricci curvature*, respectively, *deg(v)* is *in* and *out-degree* and *F(e_v_)* is the Forman curvature of edge *e_v_*, and *e_v_ ~ v* represents the set of edges incoming or outgoing on that node *v*. Subsequently, total flow on node *v* was obtained as follows:

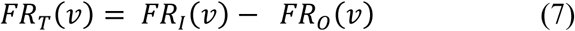

#### 2.3.3 Flux Balance Analysis

##### 2.3.3.1. Reconstruction of the stoichiometric model

For the reconstruction of stoichiometric model COBRA Toolbox (COnstraint-Based Reconstruction and Analysis Toolbox) [80, 81] was used. Necessary exchange reactions and transport reactions for uptake and excretion of typical substrates and products, respectively, were introduced into the model to avoid any metabolic gap. The model also consisted of ATP consumption reaction for its maintenance inside the model [67]. The network was devoid of any organism specific internal compartments and only external and internal environment were taken into account. Stoichiometric model is represented in the form of stoichiometric matrix, S, that is a mathematical representation of the network. In this matrix, each reaction is a column and each metabolite is a row. And the participation of the metabolites in the corresponding reactions is denoted by their stoichiometric coefficients where ‘-1’ is used for the consumption and ‘+1’ for the production of the metabolite. In our model, the upper and lower bounds of reversible reactions were set between +1000 and -1000, and for irreversible reaction between 0 to +1000. Uptake and excretion reactions were also set between -1000 and +1000.

##### 2.3.3.2. FBA simulation

Once the model is reconstructed, constraint-based FBA simulation was performed in COBRA Toolbox [80, 81] in MATLAB [82]. FBA assumes steady-state kinetics and uses linear programming (LP) based optimization to determine the flux distribution to solve a given metabolic objective function by maximizing or minimizing it [83]. Hence, the LP problem was formulated as,

Maximize Z,

Subject to

*S ● v* = 0

*v*_min_ ≤ *v* ≤ *v*_max_

Where, Z is the objective function to be maximized (or minimized), S is the stoichiometric matrix of *m* x *n*, *v* represents the flux vector that is controlled through enzyme capacity constraints *v_min_* and *v_max_* representing lower and upper bounds, respectively. After conversion into mathematical form, the simulation was performed to maximize the objective function. Initially the objective function was carrying zero flux indicating metabolic gaps in the model. The required demand reactions were added to the network to refine the model till the non-zero value of the objective function.

## 3. Results

### 3.1. Model Reconstruction and Network topology

Initially, after editing the *Leishmania major* metabolic network **iAC560**, as mentioned in section 2.1, a directed master network, **M**, was built. This model was used to construct a bipartite directed graph as follows: all transport and exchange reactions were removed; all self-loops, currency metabolites (ATP, ADP, CO_2_, etc.), electron carriers (NADP^+^, NADPH, NAD, NADH), Waters (H_2_O), and electron transporters (H^+^, PO_4_^+3^, HCO_3_^-^) were removed; each reversible reaction in the network was converted to two irreversible backward and forward reactions; then, each irreversible reaction was considered as directed edge connecting the substrate metabolite to the reaction or reaction to the product metabolite; duplicate reactions and metabolites were carefully observed in the network and removed. Moreover, each directed edge was assigned with a weight corresponding to the stoichiometry of the involved metabolite (substrate or product) in the reaction under consideration. This model consisted of 226 metabolites (Table1, Figure 3).

**Figure 3.**
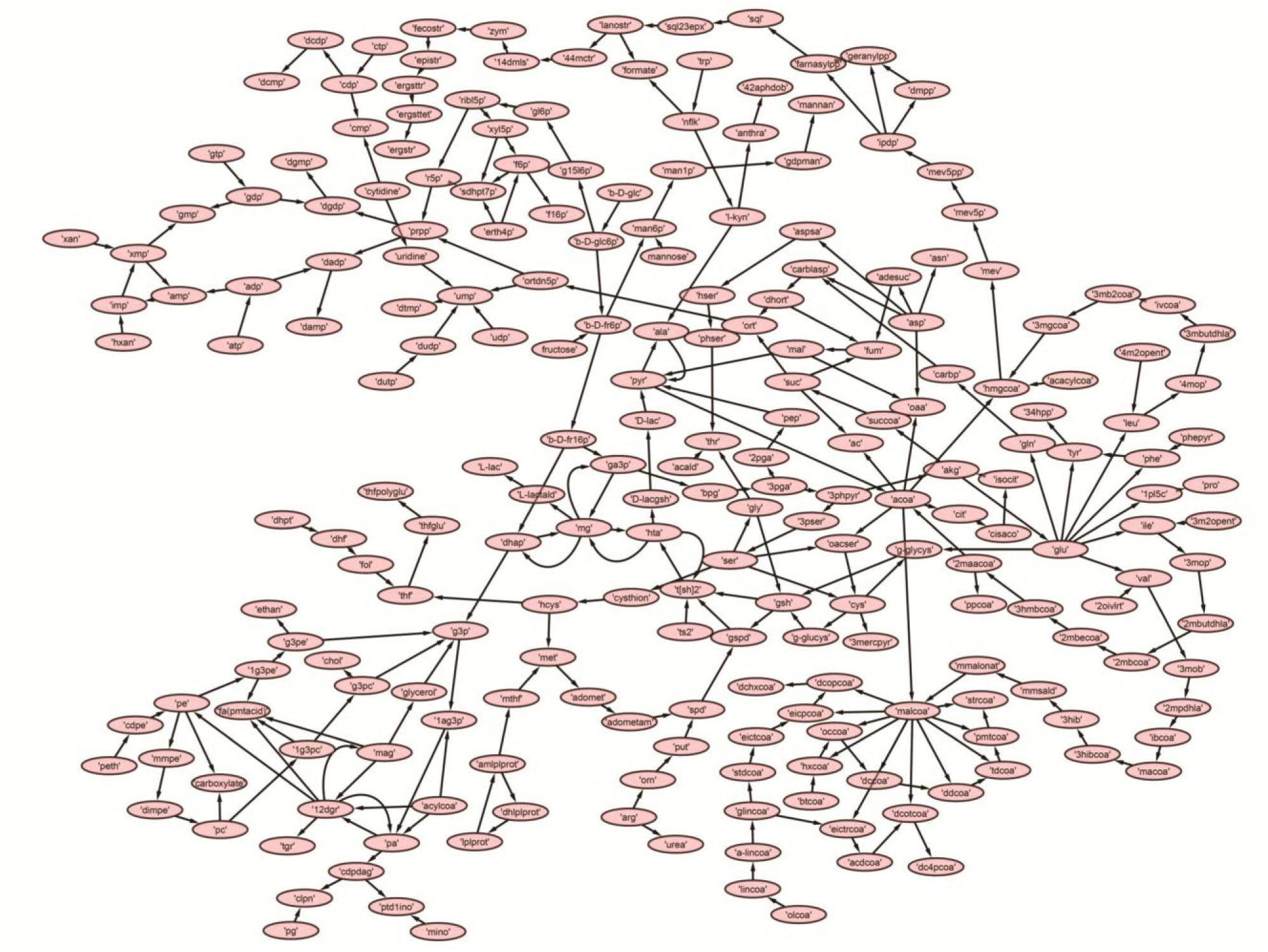
The reconstructed master model, M, after removing currency metabolites and self-loops

Further, to get a deeper insight of the *Leishmanial* metabolic network, the master model, **M,** was subdivided into 7 small directed bipartite graphs (M1, M2, M3, M4, M5, M6, and M7) (Table 1, Figure S1). Briefly, M1 consisting of 39 components was built by including glycolysis, mannan synthesis, TCA cycle and PPP pathway. M2 comprised of 81 species from amino acid metabolism, M3 (23 metabolites) has fatty acid synthesis and degradation, M4 (23 metabolites) represented sterol biosynthesis. In M5, Glycerolipid and glycerophospholipid metabolism (27 components) were included. Nucleic acid metabolism with 36 metabolites was presented in M6, and for M7, methylglyoxal (MG) and trypanothione (T[SH]_2_) metabolism (33 metabolites) were chosen.

**Table 1.** Topological properties of the Networks

| **Model** | **No. of metabolites** | **Clustering Coefficient** | **Network diameter** |
| --- | --- | --- | --- |
| M | 226 | 0.038 | 31 |
| M1 | 39 | 0.026 | 17 |
| M2 | 81 | 0.022 | 13 |
| M3 | 23 | 0.118 | 8 |
| M4 | 23 | 0.051 | 17 |
| M5 | 27 | 0.046 | 12 |
| M6 | 36 | 0.017 | 8 |
| M7 | 33 | 0.010 | 12 |

The network topology of all networks was analyzed in Cytoscape and is summarized in Table 1 and Table S1. The clustering coefficient of the master network, M, was 0.038. Among the models M1-M7, only M7 had the lowest clustering coefficient 0.01 (Table 1). The range for clustering coefficient lies between 0-1 where ‘0’ denotes no interconnectivity and ‘1’ represents highly interconnected network. The clustering coefficient of M7 indicates that the network has lesser connectivity in comparison to other networks, showing the transmission of the signal is linear and in one direction.

Further, betweenness centrality calculated for each node in M network pointed out important metabolites. Nodes having high betweenness centrality are known to act as bridge between nodes for transmission of information in the network. In our M model several of these metabolites (HTA, MG, Pyr, T[SH]_2_) with high betweenness centrality were part of M7 model (Table S1) suggesting the importance of M7 model.

### 3.2. Curvature distribution in master network

Normalized (Forman) and unnormalized (Forman-Ricci) Forman curvatures of directed edges and nodes were calculated in all reconstructed models according to the section 2.3.2. For each vertex in all directed graphs, we computed Forman curvature (Fv), Out Forman (Fo), In Forman (Fi), Out Forman-Ricci (FRo), In Forman-Ricci (FRi) curvatures, total Forman and Forman-Ricci flow.

Forman curvature of nodes and edges in network ‘M’ was computed and their distribution was plotted to analyze their frequency (Figure 4). From the plots, it was observed that most of the nodes and edges were comprised of negative curvature values. Although, the distribution of Forman curvature of both nodes and edges was broad, the peaks with high frequency were concentrated towards (or at) ‘zero’. The distribution of Fi and Fo of nodes were found to be slightly different than Forman curvature of nodes however the peaks between 0 and -2 were found to be with frequency greater than 50. Similar was the case with Fi and Fo of edges where one peak in both the plots had frequency greater than 50. Further extraction of these peaks demonstrated that metabolites belonging to amino acid metabolism, sterol synthesis, fatty acid biosynthesis and degradation, glycerolipid and glycerophospholipid metabolism pathways were part of these high peaks. Interestingly, peaks appearing far from ‘zero’, those with higher Forman curvature, consisted of edges and nodes related to glycolysis, MG metabolism and T[SH]_2_ metabolism pathways. Furthermore, these metabolites were also spotted as important vertices from Forman curvature plot of nodes in ‘M’ model (Figure 5).

**Figure 4.**
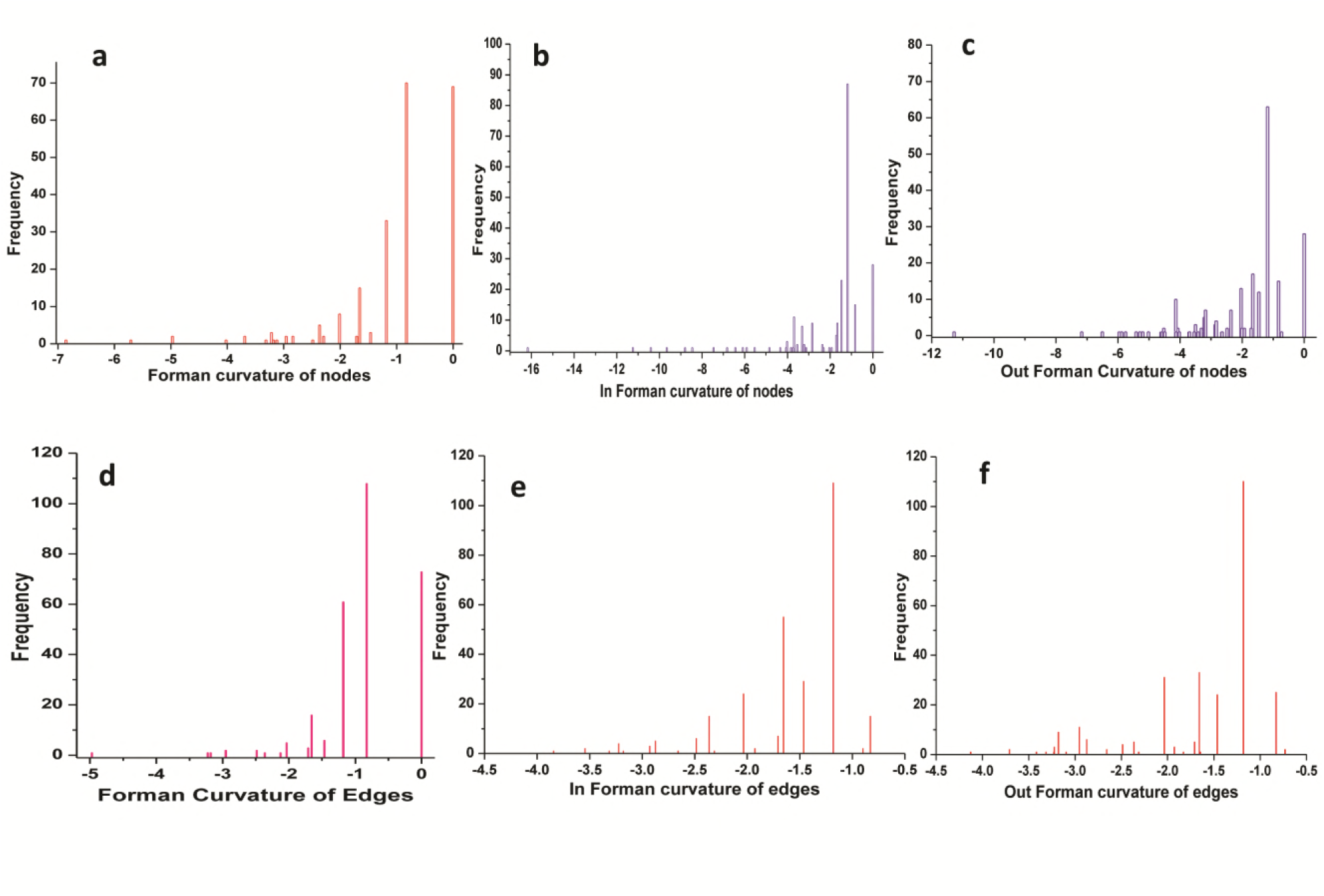
Distribution of Forman curvature of directed edges and nodes in master (M) network.

**Figure 5.**
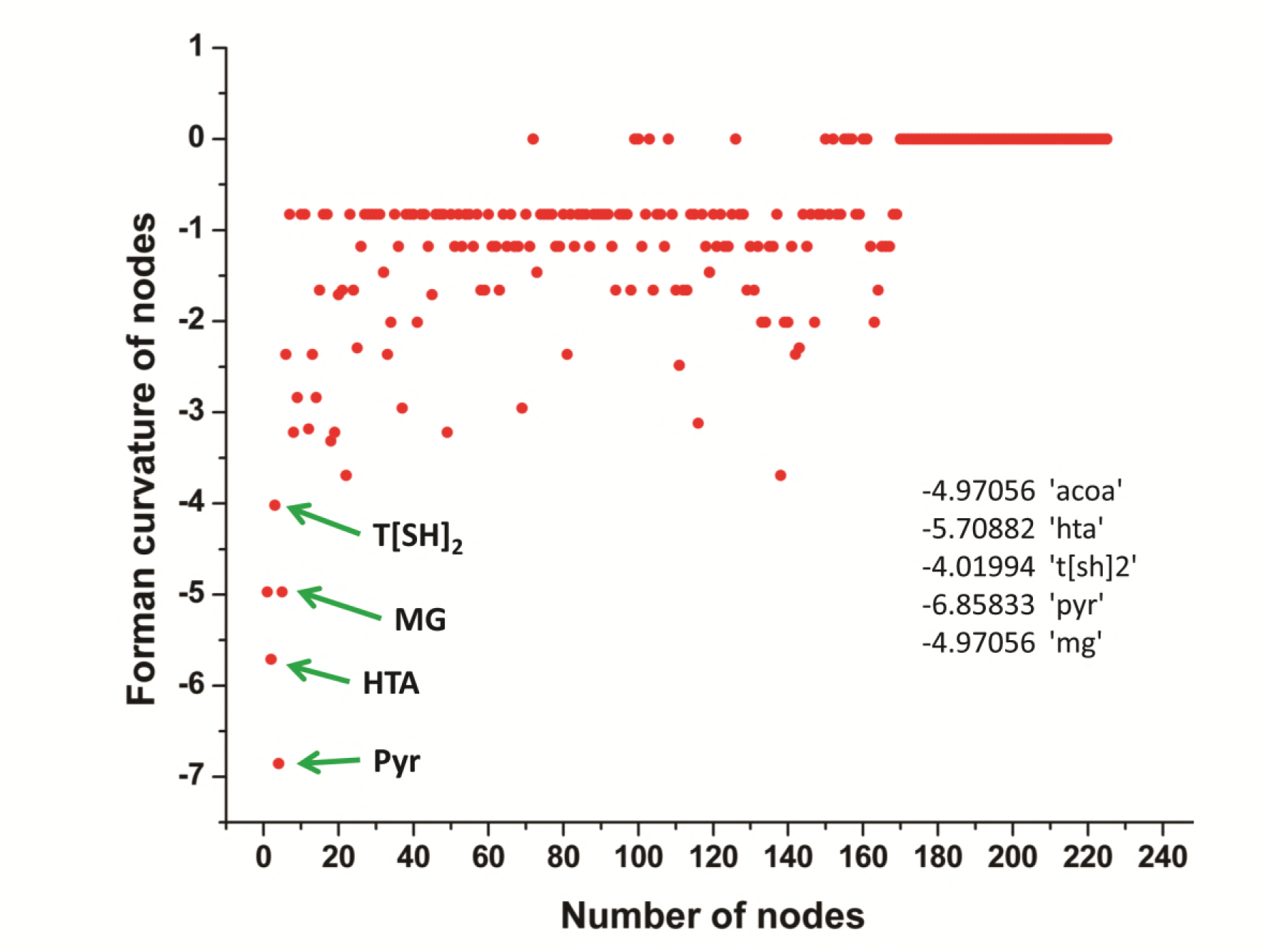
Representation of Forman curvature of nodes in master model. Metabolites with high curvature values are labeled and pointed with green arrows.

### 3.3. Correlation between Forman curvature and common network measures

Various common statistical topological measures were calculated for all networks and studied to find any significant correlation with the curvature measures (Table 2).

**Table 2:**
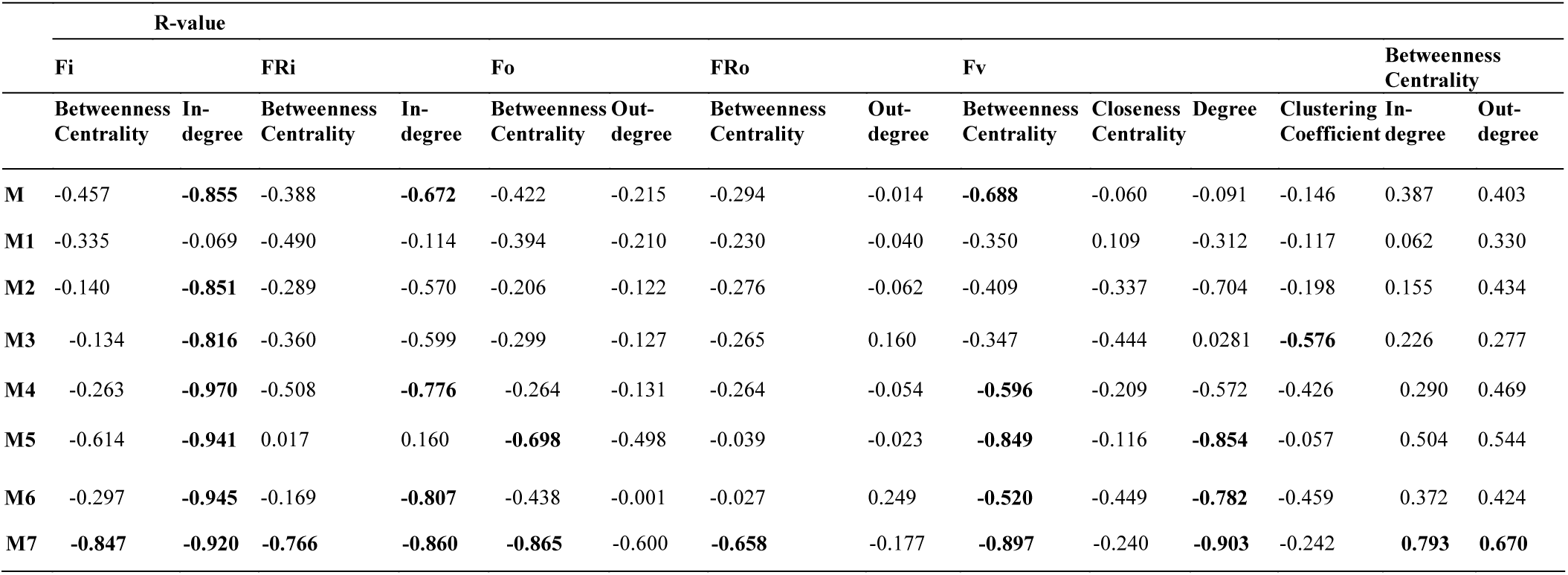
Summary of R-values for different models.

#### 3.3.1. Master model

The correlation between Forman curvatures and betweenness centrality, closeness centrality, In-degree and Out-degree were computed in ‘M’ model (Table 2, Figure 6). A high negative correlation was observed for Fi and FRi with and In-degree (Figure 6b, 6f). Moderate negative correlation was reported for Fi, FRi, Fo with betweenness centrality (Figure 6a, 6c, 6e). Interestingly, In-degree and Out-degree have also shown moderate correlation with betweenness centrality but the R-value was shifted to positive side than the negative one (Table 2). Moreover, in contrast, Fo and FRo have shown very less or near negligible correlation with Out-degree and betweenness centrality, respectively (Figure 6d, 6g, 6h). Forman curvature of nodes was also negatively correlated with high magnitude to betweenness centrality than that of closeness centrality suggesting the importance of the former in finding important nodes in the network (Table 2).

**Figure 6.**
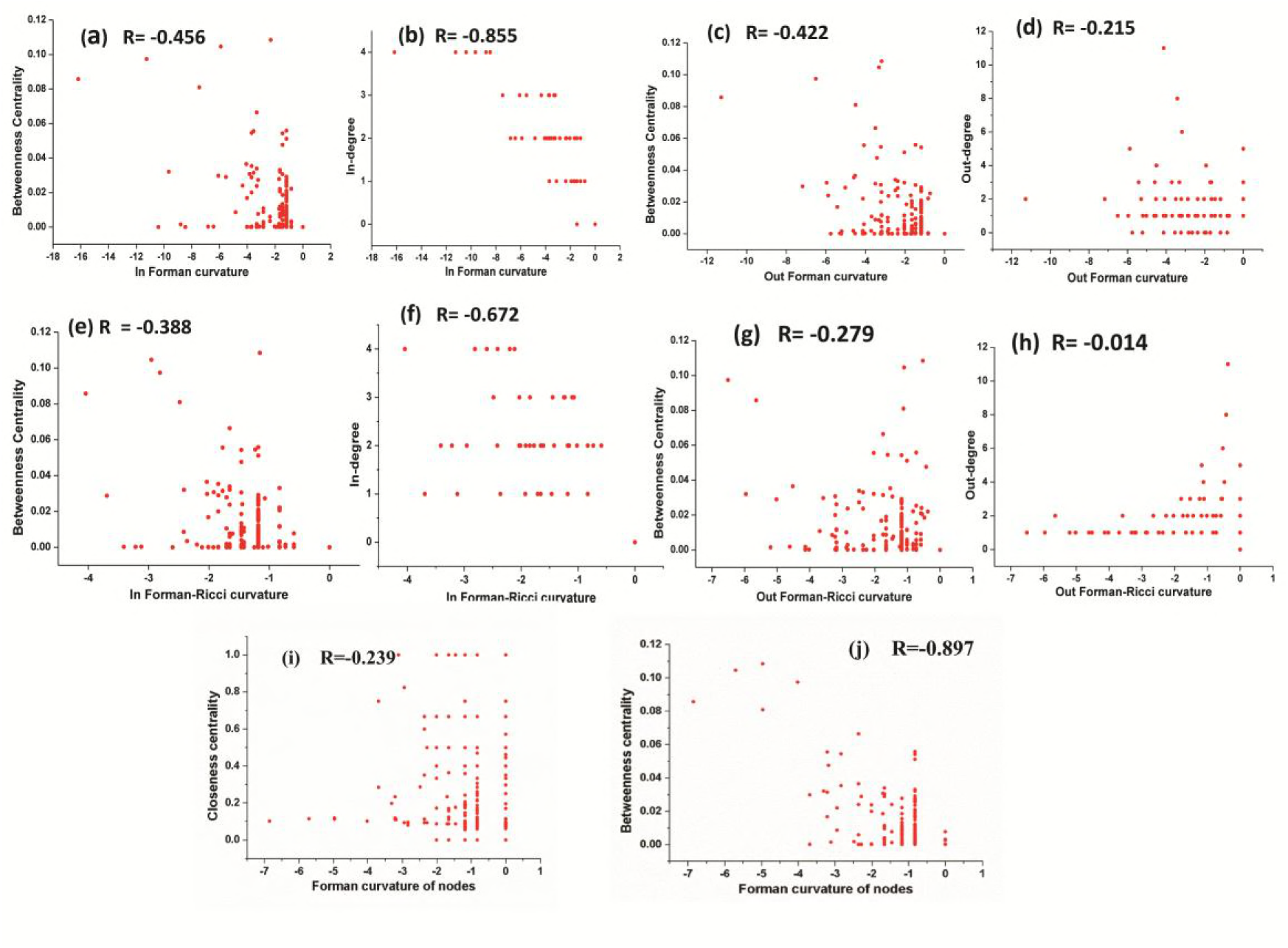
Correlation between (a) Fi-Betweenness centrality; (b) Fi-In-degree; (c) Fo-Betweenness centrality; (d) Fo-Out-degree; (e) FRi-Betweenness centrality; (f) FRi-In-degree; (g) FRo-Betweenness centrality; (h) FRo-Out-degree; (i) Fv-Closeness Centrality; and (j) Fv-Betweenness Centrality of nodes in Master ‘M’ network. Also, Spearman correlation coefficient is indicated for each.

Importance of nodes can also be identified by employing pagerank [84, 85]. Thus, we calculated correlation for pagerank with Forman and Forman-Ricci curvature in ‘M’ model. Our model showed that the correlation for pagerank with Fi and FRi was observed high negative than Fo and FRo. Interastingly, similar high but positive correlation was found between pagerank and in-degree but with out-degree it was noted very less.

#### 3.3.2. Small scale models

A clustering coefficient is the measurement of the degree to which nodes in a network tend to cluster together, thus, it is said that clustering coefficient is an important measure to quantify the curvature of complex networks [28, 86]. Moreover, *degree* of a node represents the total number of edges connected on a node. Therefore, we investigated the correlation of Forman curvature of nodes with degree and clustering coefficient. It was noted that, in all models (M1-M7) except M3 (-0.576), the value of correlation between Forman curvature of nodes and clustering coefficient was very weak negative in comparison to correlation between Forman curvature and degree. Interestingly, the highest negative correlation between Forman curvature and degree was mainly noted for model M5, M6 and M7 (-0.854, -0.782, and -0.903, respectively) (Table 2). It might be possible because of the nature of Forman curvature that captures the geometric properties of classical notions discretized by networks.

Betweenness centrality is another important measure of network topology. This measures the degree of a node passing through the shortest path to reach other nodes [5]. Similarly, closeness centrality [87] of a node measures the mean distance from a vertex to other vertices and represents the degree of closeness of nodes. In terms of correlation, we obtained weak negative correlation coefficient between closeness centrality and Forman curvature of nodes. M1 model showed positive correlation but it was very weak to be considered. In contrast, in most of the models (M4-M7) along with the ‘M’ model, Forman curvature was found to be negatively highly correlated with betweenness centrality (Table 2).

We find a high negative correlation for Fi and In FRi with In-degree in all models. On the other hand, a good negative correlation of betweenness centrality with Fi and FRi was observed only in case of M5, M7 and M4, M7, respectively. Surprisingly, in contrast to Fi and FRi, correlation for Fo and FRo with out-degree and betweenness centrality was noted very weak to moderate negative. Among the small scale models, only M7 indicated high negative correlation for Fo and FRo with betweenness centrality. It was found similar to the correlation for betweenness centrality and In-and Out-degree for M7 model (high positive correlation). However, other models only showed moderately positive correlation for the same (Table 2). It was expected because the Fi and FRi of a node is dependent on the edges incident on that node. However, weak negative correlation for Fo and FRo with out-degree and betweenness centrality was not expected [40].

### 3.4. Curvature and topological robustness of models

Robustness of the model was investigated on the basis of the Forman and Forman-Ricci curvature of the nodes. For this, we computed these curvatures across all edges and nodes of the network and observed that the curvature for all networks (M-M7) is negative (Figure 6, Table S1). High curvature nodes are said to be the *backbone* of the network those act as *bridges* between major network communities [89] (Figure 7, Figure S1). Since, the nodes/edges with higher curvature values act as *backbone* of the network [89], we wanted to look for the overall effect of these high curvature nodes/edges on the network robustness. For investigating the robustness of the model, the nodes were removed on the basis of decreasing strength in terms of Forman curvature, Forman-Ricci curvature and clustering coefficient. Nodes with low curvature and low clustering coefficient were removed first. The robustness of the network was measured in terms of ‘network connectivity’.

**Figure 7.**
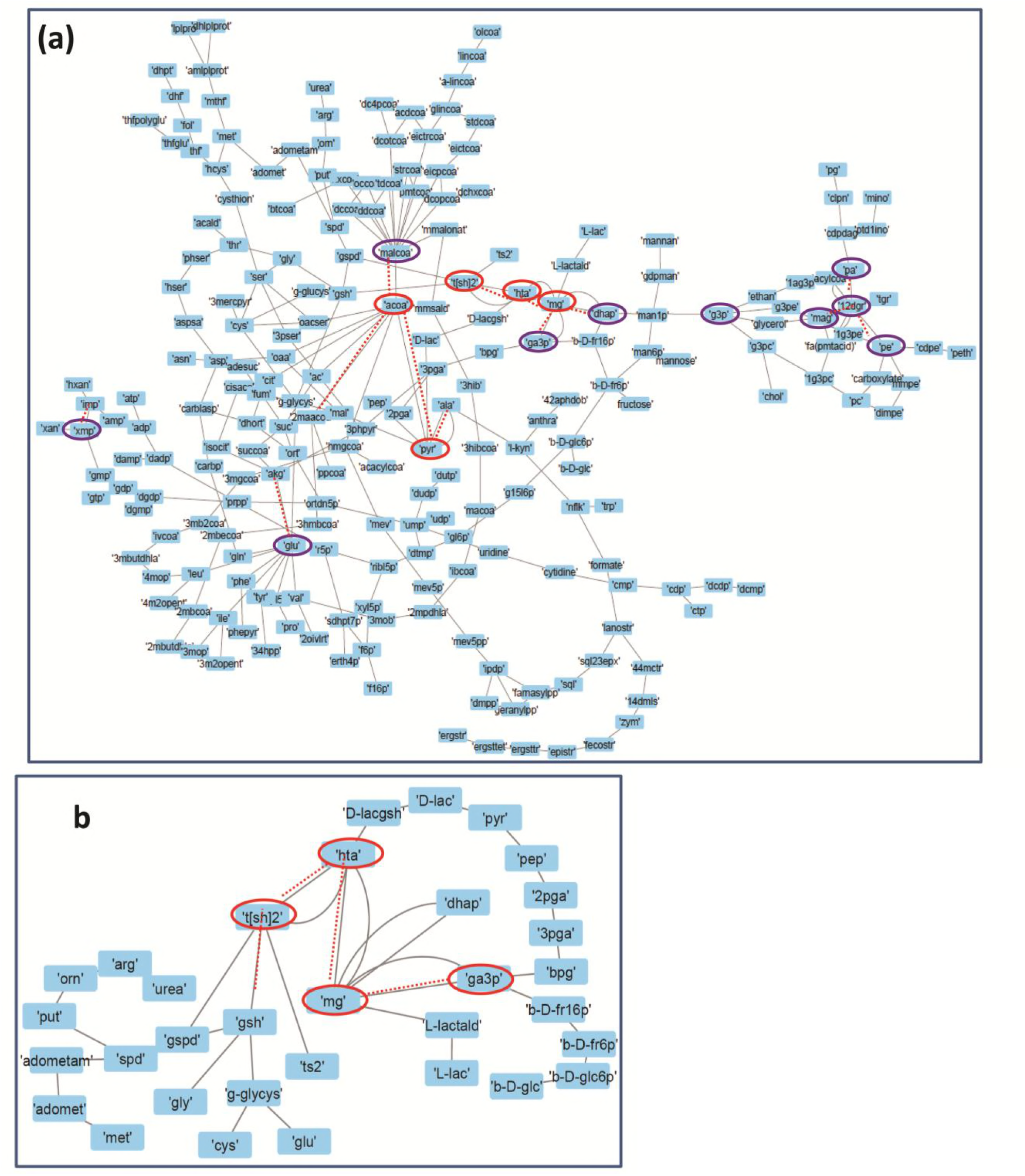
Depicting the internal connectivity in (a) Master model, and (b) M7 model. The components are connected by *backbone* forming high curvature nodes (encircled in red color). Low curvature nodes are shown without encircled nodes. High curvature nodes act as *bridges* between major network communities and play important role in determining structural and functional organization of network.

Connected components of a network tell us about the connectivity of the nodes in the network. Highly connected network is represented as ‘1’, and a higher value is an indication of disconnected network. In Figure 8, we plotted network connectivity against the nodes removed on the basis of Forman curvature and clustering coefficient. It was observed from the plot that removal of high curvature nodes caused faster disintegration of the network in comparison to low curvature nodes. Nodes deletion according to Forman-Ricci curvature gave similar result as with Forman curvature. However, nodes removal on the basis of clustering coefficient has also led to the network destruction but the process was found to be slow when compared with nodes deletion on the basis of Forman curvature. Also the level of disintegration of the network was found at lower rate in the former case (Figure 8).

**Figure 8.**
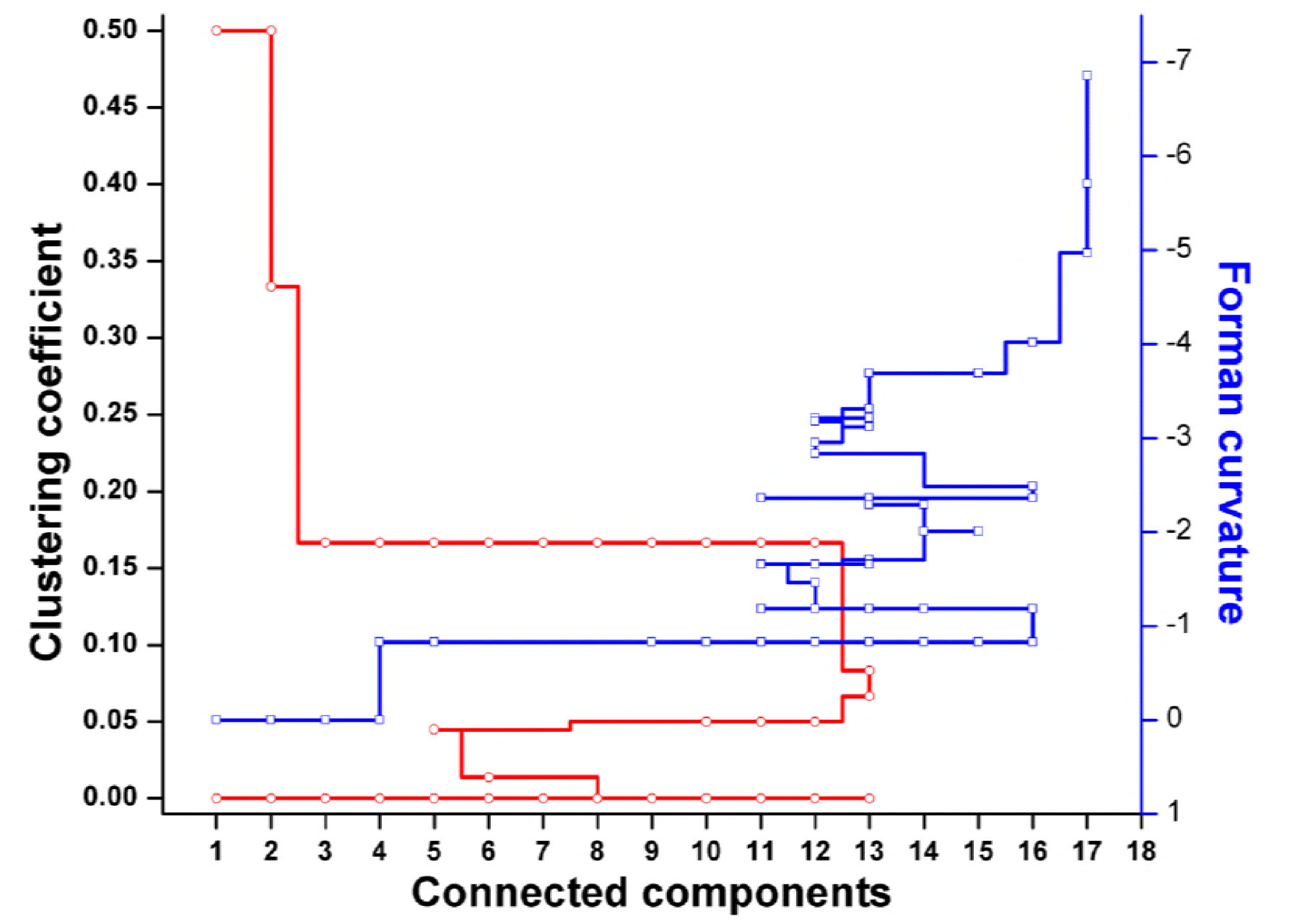
Robustness analysis of the Master model (M) using nodes deletion on the basis of Forman and Forman-Ricci curvature, or clustering coefficient.

In our master model we found total 25 nodes with high curvature values (above -2.0), among them 13 were with curvature greater than -3.0 (Table S1). We noted that most of the nodes from these were related to ‘M7’ model that consists of glycolysis, T[SH]_2_ synthesis, and methylglyoxal metabolism pathways, pointing out the importance of ‘M7’ model.

### 3.5. Flux balance analysis

#### 3.5.1. Characteristics of the reconstructed network

As stated in previous section, the role of several metabolites including MGO and T[SH]_2_ via Forman curvature and Forman-Ricci curvature measures was found important for maintaining the integrity of the master model. In our small networks, these metabolites belong to model ‘M7’, hence we manually reconstructed this model for FBA. Since our focus for performing FBA was to investigate the importance of glyoxalase pathway in redox metabolism, we refrained our model from genome-based construction and kept it at simple metabolic level. The proposed stoichiometric model accounts for the reactions of glycolysis, glyoxalase pathway, and thiol metabolism. The reconstructed model resulted in a network consisted of 109 reactions that included 48 main reactions, 22 transport reactions and 24 exchange reactions (Table 3). ATP maintenance and NADPH demand reactions were also included to fulfill their need in the system. The exchange reactions were added to allow input-output exchange of the extracellular metabolites to enter the system from the medium or excretion of end products of any reaction out of the system. All reactions were set to the maximum for upper and lower bound limits for reversible/irreversible, uptake/excretion reactions except for glucose [90]. For this small scale model we took utmost care in including only the reactions those are required for maintaining the production of desired metabolites. Substrate uptake and product excretion reactions were added only if required.

**Table 3.** Properties of FBA model constructed from M7 model

| Property | Count |
| --- | --- |
| Reactions | 48 |
| Transport reactions | 22 |
| Exchange reactions | 24 |
| Metabolites | 86 |
| Compartments | 1 |

##### 3.5.1. Formulation of objective function

Biomass objective function (BOF) is a mathematical equation that provides drain (demand) of essential metabolites needed to mimic the cellular growth [91, 92] by solving the linear programming problem using FBA. For our model, we integrated the objective function consisting of the pseudo consumption of precursors from different pathways present in the model. Our aim was to observe the effect of metabolism of MGO through Glyoxalase pathway on the system; hence, we focused on the formulation of objective functions (OF) by taking into account the metabolites only used in the model:

***Objective function: pyr + D-lac + atp + h2o -> adp + pi + h***

The end product of glyoxalase pathway is D-lactate that ultimately leads to the formation of pyruvate. When MGO is in excess, as mentioned earlier, it can cause the formation of free radicals. Hence, separate objective functions were designed in such a way that they shall give an idea of how MGO will control our model system and to what extent. For simplicity, the stoichiometric coefficients of the precursors and metabolites used in formulation were kept as default. We considered, for our model, the end products of both the pathways, glycolysis (pyruvate) and glyoxalase pathway (D-lactate), contributing to the total biomass production. Hence, the system was investigated in two scenarios, first, functional GLOI (Glyoxalase I), and second, nonfunctional (absent from the system) GLOI. Glucose was supplied as carbon source in both the scenarios.

###### 3.5.1.1. Scenario 1: functional GLOI

First scenario corresponds to functional GLOI by keeping its flux higher. It was noticed that glycolysis carried constant flux to produce pyruvate. However, glyoxalase pathway, comparatively, carried maximum flux for the production of its end product D-lactate. It was also noticed that in presence of functional GLOI, advanced glycation end products (AGEs) formed at very minimal level suggesting the important role of glyoxalase pathway. Although, both the pathways carried different fluxes level, the maximization of OF was seemed to be majorly contributed through MGO synthesis and GLOI/II activities (Figure 9a). Other metabolites synthesized at very minimal level probably due to smaller scale of our model and lack of other reactions for the compensation of their consumption.

**Figure 9.**
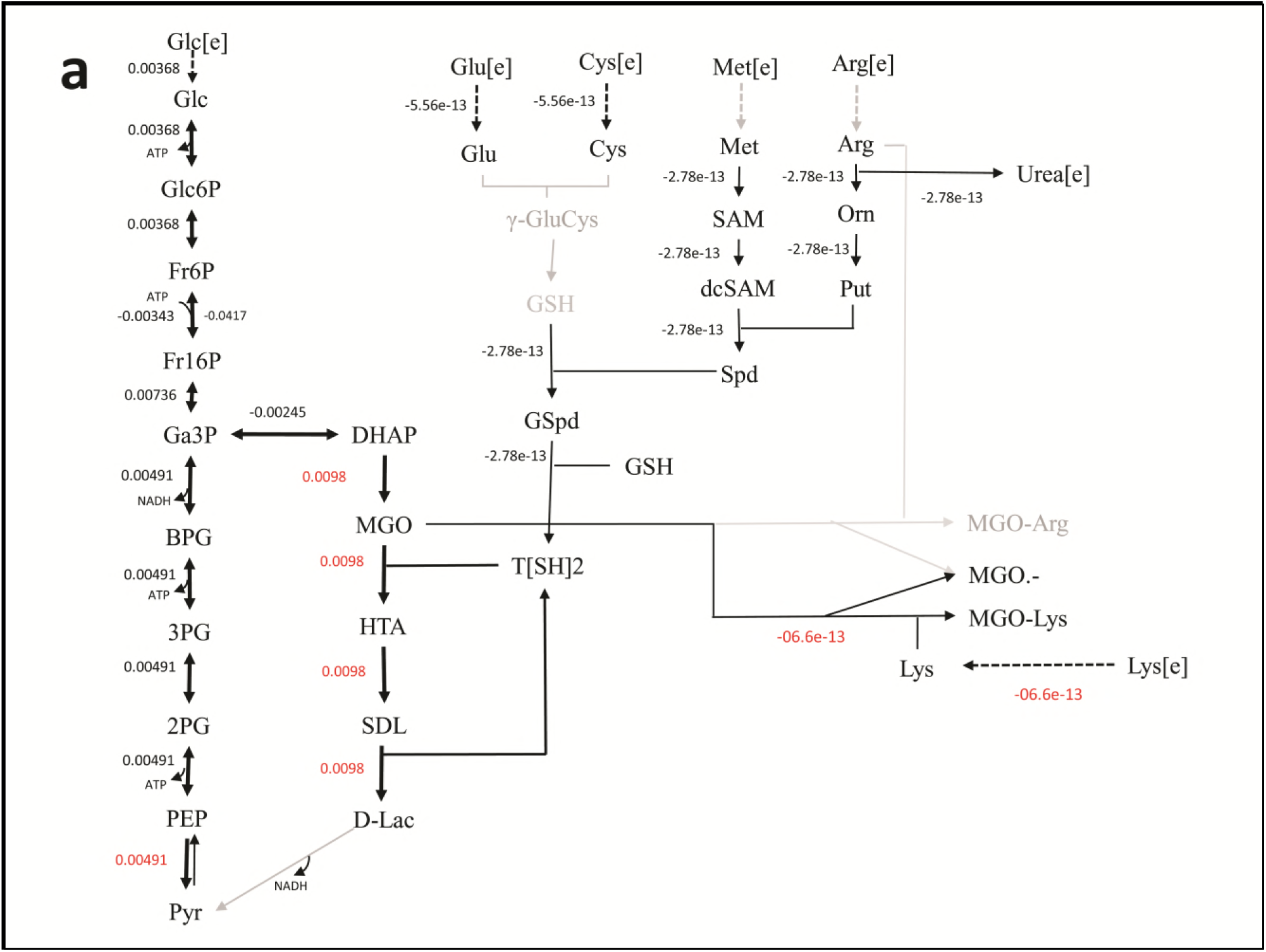
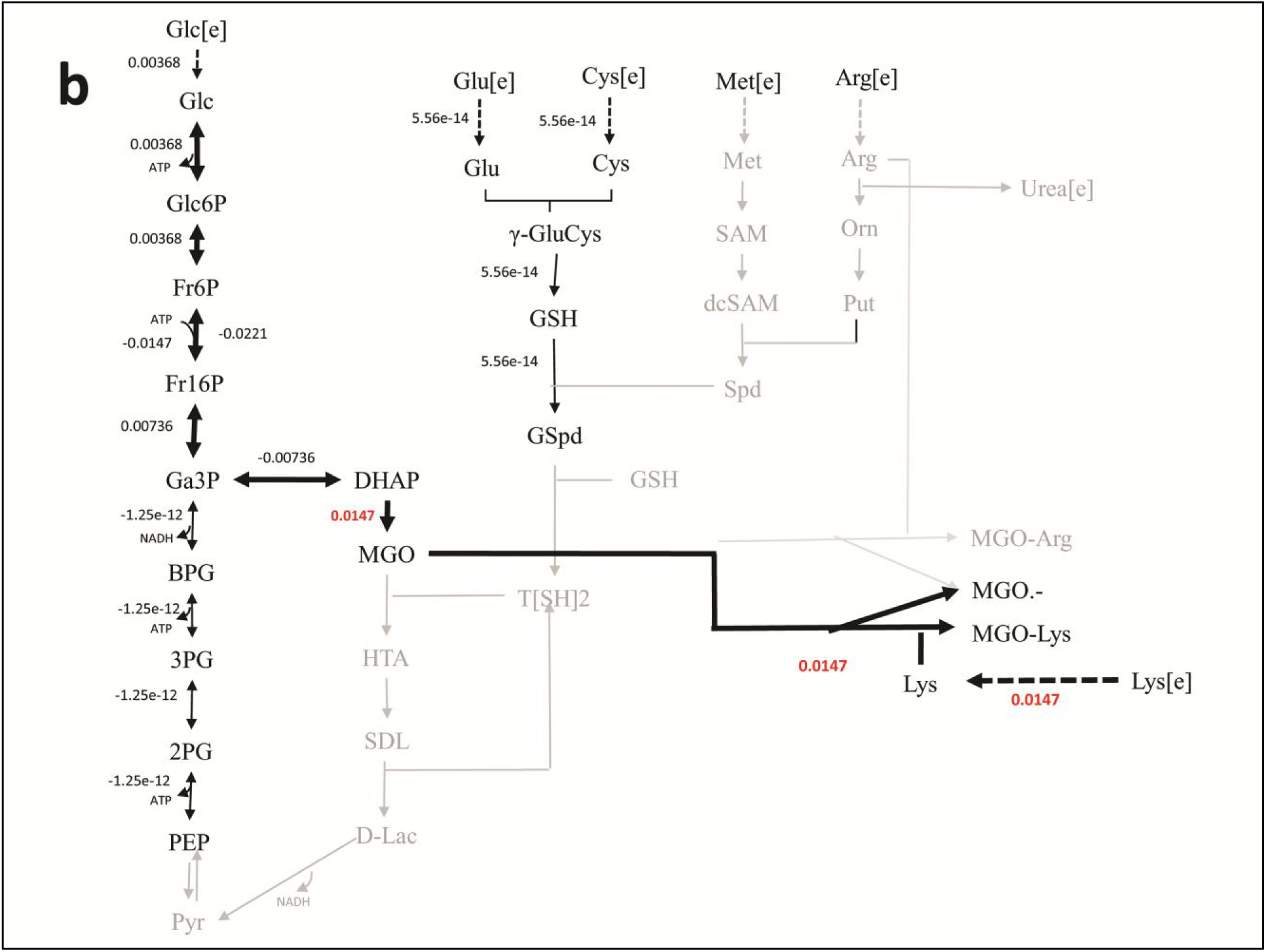
Reaction fluxes for **(a)** scenario 1 (Functional GLOI), and **(b)** scenario 2 (Non-functional GLOI). The depicted scenarios are illustrating the flow of flux for the objective function (OF). The simulation was performed for the production of pyruvate (from glycolysis and glyoxalase pathway) and role of MGO in producing AGEs (MGO-Arg and MGO-Lys) and MGO●-free radical. Solid thick black arrows represent the direction of main flux flow through the model. Gray arrows are an indication of undetermined (zero) flux through the respective reaction. Dotted arrows represent the transport of the metabolites.

###### 3.5.1.2. Scenario 2: non-functional GLOI

In, second scenario, we blocked the flux through glyoxalase pathway by setting GLOI flux to ‘zero’ that makes it inactive in the pathway. When the simulation ran in this case, we noticed that glucose entered the system at the same rate as in scenario 1. Even the flux rate was found to be more or less similar for glycolysis but only upto the synthesis of GA3P and DHAP. In fact, the flux through TPI enzyme (DHAP synthesis) was more than that of scenario 1 that probably contributed to the synthesis of MGO. It is noticeable that MGO was produced at twice the rate than the previous case. This higher level of accumulation of MGO was found to contribute to equal level of the formation of AGEs and MGO^●-^ free radicals (Figure 9b). It was worth noticing that in case of inactive GLOI, the flux moved from glycolysis to for the synthesis of MGO. In the model, this resulted into negligible flux through the rest of the glycolysis pathway. Although in real network, pyruvate production will be happening at normal rate except for that the inactivation of GLOI should lead to the increased level (~1.5 fold) of accumulation of MGO. It was observed that, inactivation of GLOI caused the ‘zero’ flux through the rest of the glyoxalase pathway.

## 4. Discussion

Due to the complex nature of biological networks it has been a major challenge to unravel their architecture and decipher the structure-function relationship. To discover the robustness and reliability of a biological network it is important to understand its basic structural framework by quantitatively analyzing through mathematical tools [4-8, 93,94]. This can help in uncovering the meaningful information by reducing the system to a smaller level in order to understand the relationship among its various components [9-26]. In this paper, we motivate the use of systems biology driven approaches including *Forman curvature* and *Forman-Ricci curvature* measures [95, 96] and their associated geometric flow along with the FBA for the analysis of complex metabolic network of *Leishmania* parasite.

Through curvature measures and topological analyses we were able to understand the structural and functional relationship among the various metabolites in our network. Forman and Forman-Ricci curvature of the master ‘M’ network has shown that nodes/edges with high curvature values differ drastically in structure from the ones with low curvature values. The ones with high value appear to be in denser clusters and acted as bridges between the major sub-clusters in the network. This measure not only helped us to identify the relatively more important parts of the network but also provided the information on identifying the sub-networks. This gave us the idea to break the network in sub-models to study the properties of various pathways at smaller scale and has helped us to detect important substructures inside the network belonging to particular classes of vertices or edges. Importantly, our analyses have pointed out several central nodes belonging to very common yet crucial metabolic pathways those include glycolysis, glyoxalase pathway, and T[SH]_2_ metabolism pathways (Figure 5, 7, S1). The significant importance of these high curvature nodes (pyruvate, acetyl coenzyme A, methylglyoxal, hemithioacetal, trypanothione) has also been demonstrated while performing the robustness analysis on the network (Figure 8). In fact, high correlation between Forman/Forman-Ricci curvature measures and common network properties has also demonstrated and pointed out the structural and functional importance of ‘M7’ model that comprises of above mentioned metabolites (Table 2).

It is worth noticing that several of these metabolites (MGO and T[SH]_2_) have very important role in the regulation of redox homeostasis of the parasite. As mentioned earlier, MGO is highly cytotoxic and mutagenic carbonyl species that can react with DNA and various proteins and enzymes, causing their irreversible modification and inactivation [97-103] that leads to the generation of advanced glycation end products (AGEs), cross-linked adducts and MGO^●-^ free radicals. Hence, the most needed elimination of MGO is carried out by an ubiquitous thiol dependent glyoxalase system that comprises of two key enzymes, glyoxalase I (GLOI) and glyoxalase II (GLOII) [104, 105]. These two enzymes sequentially convert MGO into D-lactate in a two steps process: spontaneously formed MGO-T[SH]_2_ conjugate, hemithioacetal (HTA), is first isomerized to S-D-lactoyltrypanothione (SDL) catalyzed by GLOI, followed by GLOII mediated D-lactate formation and release of free T[SH]_2_ [106]. This D-lactate is then converted to pyruvate with the help of D-lactate dehydrogenase (D-LDH) enzyme [107, 108]. In glyoxalase pathway, GLOI is reported to be the most crucial part of the pathway and the conversion of HTA to SDL is considered the rate limiting step. The essential role of GLOI has been demonstrated earlier in GLOI attenuated *Leishmania donovani* strain [109].

Our FBA result has also shown similar observations when we simulated the model in two different conditions. Although, the OF we designed in our paper only serves as a tool to identify feasible flux across the model system, using this OF, the two different conditions applied on the model has demonstrated the key role of GLOI. The nonfunctional GLOI has caused ‘zero’ flux into the OF and led to ~ 1.5 fold increased production of MGO when compared to the wild type scenario with functional GLOI (Figure 9a and 9b). The increased production of MGO in the system caused its accumulation due to underdetermined flux in the downstream glyoxalase pathway and, hence, lack of its detoxification. Among the amino acids, MGO mainly reacts with arginine and lysine, hence, the AGEs production was introduced in the form of MGO modified arginine and lysine residues. The accumulation of MGO in the system consequently led to the formation of AGEs and MGO^●-^ free radicals. Moreover, in scenario 2, glyoxalase pathway has also reflected the rate limiting step in very agreeable manner in the form of carrying the ‘zero’ flux through the glyoxalase pathway reactions. These findings revealed the importance of GLOI and are in the agreement with the previous reports. Although, due to maintaining the simplicity of the model, we have not shown the effect of MGO^●-^ free radical on other components in the system, however it has been demonstrated earlier that the excess amount of MGO causes the formation of increased level of AGEs and MGO^●-^ free radicals [110-116] and can inhibit the growth of *Leishmaia* [109].

Having stated above, for most part of the reconstructed network, taking into account the different point of observations, interaction dynamics does not drastically changed. Nonetheless, there might be instances at which major changes can occur in the network often in response to the metabolic or environmental stress. On the other hand, it might also be postulated that metabolic rewiring is driven from internal or external stress, thus causing a change in the functional requirement of the metabolites. Optimization based strategies introduced in terms of the objective function help in the inference of varied kind of interaction which ultimately characterizes the activity of individual metabolites and/or predicts their interactions with other metabolites, including unknown predictions. These unknown predictions may serve as targets for experimental validation. Thus, the interaction dynamics of GLOI as a target could be deciphered with the numerical evaluation in two scenarios (functional and nonfunctional GLOI). This further suggests that the biological parameters chosen with varying combination of parameters is not redundant. The application of certain specific combinations has helped us to avoid the lack of convergence in the reaction flux model. Furthermore, this has helped us to rule out the absence of any unique set of parameters minimizing the occurrence of noise in the said model system. The integrated analysis laid in this paper from computational perspectives helped pinpoint an interesting phenomenon that the underlined network model structure is largely preserved across the large common substructure models.

## 5. Conclusion

In the quest to improve the current understanding of biological networks we have devised a newer strategy of formulation of curvature measure along with FBA in order to explain and predict biological targets. Computational model discerning metabolic entity and associated phenomenon makes it possible to find models with particular model components. The methodology adopted in this paper has enabled us to determine the implicit relationship between metabolites. We envision our model developed as an inevitable integration arising from a common framework for representing metabolic model. The model prototype developed in current paper largely depends on its structure and topological components. On a broader note, we have developed our mathematical model in an iterative approach to optimize the flux of the metabolic system under limitation of certain enzymatic reactions thus leveraging the unifying framework for curvature measures along with FBA. GLOI deciphered as important target both from curvature as well as FBA enabled our deeper understanding into the parasite redox metabolism.

